# SOCIAL INFORMATION AND BIOLOGICAL INVASIONS: CHORUS SONGS ARE MORE ATTRACTIVE TO EUROPEAN STARLINGS IN MORE RECENTLY ESTABLISHED POPULATIONS

**DOI:** 10.1101/2021.07.24.453665

**Authors:** Alexandra Rodriguez, Martine Hausberger, Laurence Henry, Philippe Clergeau

## Abstract

When biological invasions by animals occur, the individuals arriving in novel environments can be confronted with unpredictable or unfamiliar resources and may need social interaction to improve survival in the newly colonized areas. Gathering with conspecifics and using social information about their activities may reveal the location of suitable feeding and breeding sites. This could compensate for the absence of individual information about new habitats and constitute an advantage for new settlers. If a tendency to gather in response to social stimuli is transmitted from one generation to the next, and if the benefit of gathering is lower in long time established populations, there should be behavioural differences in receptivity to social cues in populations with different colonizing histories. We hypothesized that individuals of a social species like the European starling from relatively recently-established populations would be more responsive to social cues than individuals belonging to long-established populations. We conducted playback experiments using a starling chorus to test its acoustic attractiveness to populations of starlings with different colonizing histories. We compared the reaction of individuals from two populations in rural Brittany, western France, established for a long time, with three more recently settled ones: two populations from a propagation front in southern Italy and one urban population from Rennes city in Brittany. Our data supported our hypothesis: individuals from more recent populations were more responsive to the acoustic stimulus, and gave more calls in flight than individuals from populations with an older settlement history. We discuss the different behavioural responses we observed in the different populations and the potential effects of habitat characteristics and starling densities.

## RESUMEN

Los individuos que llegan a un nuevo hábitat en las primeras etapas de una invasión biológica pueden verse confrontados a una distribución impredictible de los recursos y por ello pueden necesitar recurrir a interacciones sociales para aumentar su probabilidad de supervivencia en las nuevas areas colonizadas. Agruparse con otros individuos de su misma especie y utilizar la informatión social producidad por las actividades de estos últimos puede permitir la localización de áreas ventajosas de alimento o de reproductión. La información social obtenida de esta forma puede constituir una ventaja y compensar la falta de información personal de los nuevos individuos que llegan a ese nuevo hábitat. Si existe una tendencia a agruparse al detectar señales sociales y si esta tendencia puede ser transmitida de una generatión a la otra podrían existir diferencias comportamentales en la receptividad a las señales sociales entre poblaciones cuyas historias de colonización difieren.

Emitimos una hipótesis según la cual los individuos de poblaciones recientemente establecidas en nuevas áreas geográficas serían más receptivos a señales sociales que los individuos pertenecientes a poblaciones establecidas durante un largo lapso de tiempo en un área bien conocida por la población. Para explorar esta hipótesis llevamos a cabo un estudio sobre un ave social invasora, el Estornino pinto. Para ello realizamos experimentos de difusión acústica gracias a grabaciones de un coro de estorninos con el fin de evaluar la atractividad de los cantos ejercida sobre individuos de poblaciones con historias diversas. Comparamos la reacción comportamental de individuos pertenecientes a dos poblaciones antiguas situadas en las areas rurales de Bretaña, region del oeste francés con la reacción de aves pertenecientes a tres poblaciones más recientes: dos poblaciones del frente de propagación situado al sur de Italia y una población urbana reciente ubicada en el centro de la ciudad de Rennes.

Los resultados que obtuvimos apoyan la hipótesis emitida: los individuos pertenecientes a las poblaciones más recientes fueron más reactivos al estímulo acústico y emetieron más frecuentemente vocalizaciones durante el vuelo que los individuos provenientes de poblaciones con un período mayor de asentamiento en sus áreas. Discutimos las diferentes respuestas comportamentales observadas y comentamos el posible efecto de otras variables como las características del hábitat y las densidades poblacionales.

## INTRODUCTION

Research on biological invasions by animals has recently focused on behavioural approaches (Holway and Suarez 1999). Various studies have concentrated on aspects of the social behaviour of invasive populations. Detailed studies of introduced invasive ant species such as the Argentine ant *Linepithema humile* and the Fire ant *Solenopsis invicta* (Holway et al. 1998, Suarez et al. 1999) found that introduced ants from colonies in propagation fronts formed large supercolonies and presented lower levels of intraspecific aggression than colonies from original anciently established populations. This reduction in territorial behaviour between individuals of neighbouring nests appears as one possible explanation for the establishment success of this species in the areas of introduction. Social life and interactions with conspecifics may be particularly relevant during the colonization process, as living in groups may constitute an advantage when facing predators, looking for resources etc. Animals can draw upon their own experience (Chesler 1969, Valone 1989) as well as upon social transmission (e.g. by social facilitation, local enhancement, observational learning) (Chesler 1969, Boyd and Richerson 1988, Galef 1993, Valone and Templeton 2002) in order to acquire appropriate information about their environment.

The use of social cues from conspecifics or information obtained from other species can be a mechanism allowing the establishment of new populations by local enhancement at suitable sites or local avoidance of unsuitable ones (Smith et al. 1999, Slaa et al. 2003). Some authors have started to study the use of information in biological invasion contexts by comparing native and introduced species. For example, Hazlett et al. (2003) found that two invasive crayfish species, the Spinycheek crayfish *Orconectes limosus* and the Red swamp crayfish *Procambarus clarkii*, were able to detect and react to the chemical alarm signals produced by conspecifics and by other crayfish species in the presence of predators whereas the native and non invasive species *Austropotomobius pallipes* was only sensitive to the conspecifics’ chemical signals. These results suggest that invasive species probably use a broader range of interspecific information concerning predation risk than native species. However no studies have compared in an intraspecific approach the degree of use of information between populations settled in a well known and anciently occupied environment and populations in more recently occupied sites as in propagation fronts. In these last contexts individuals may be confronted with unpredictable or sub-optimal environments (Boyd and Richerson 1988) and information about non-toxic food, and suitable breeding and feeding sites may be crucial (Galef and Laland 2005).

As different social cues can act as social aggregators, they may play an important role in detecting food resources, the presence of predators and suitable breeding sites (Gochfeld 1978). This assertion is supported by results on species foraging in groups like the house sparrow *Passer domesticus* (Elgar 1986) or the chicken *Gallus gallus domesticus* (Marler et al. 1986, Wauters and Richard-Yris 2003), where it has been shown that food calls induced flock formation or local enhancement around the individual producing the calls, and therefore aggregation at the food resource. The frequency of the food calls depends on the characteristics of the food source (dispersed or concentrated, attractive or repulsive), the physiological state of individuals (hungry or not) and is related to the amount of exploratory behaviour (Elgar 1986, Wauters and Richard-Yris 2003).

Just as acoustic signals seem to be important in the location of food resources in these cases, studies on colonial breeding behaviour have demonstrated that individuals, in particular new settlers, could use acoustic stimuli as signals indicating the presence of conspecifics in some potential breeding sites (Alatalo et al. 1982). Mountjoy and Lemon (1991) demonstrated that both male and female starlings *Sturnus vulgaris* were attracted by playback of male song near nest boxes, and Alhering et al. (2006) were able to attract Baird’s Sparrow males to suitable but socially empty breeding sites with playbacks reproducing breeding male vocalisations. In their experiment, they used as a control suitable and ecologically similar sites where they did not playback calls, and no birds were attracted to these sites. This confirmed that the acoustic signals were likely the key elements inducing site choice and not the quality of surrounding vegetation.

The European starling is an invasive bird that has become successfully established after its introduction to very different habitat types around the world, and it continues to spread to the southern regions in Spain and Italy (Motis et al. 1983, Feare 1984, Pasquali 1984). This bird has a complex social organization, where vocal interactions are essential. In this species song seems to play a major role in mate choice (Mountjoy and Lemon 1996) and song sharing reflects social affiliation (Hausberger et al. 1995, Hausberger 1997). Vocalisations may act as a group signature as individuals of a same group share a same dialect (Adret-Hausberger 1983) and can be a social “marker” that allows individuals of a same group to recognise and join each other in roosts (Hausberger et al. 2008). Acoustic social cues can thus be considered as central parts of the flocking and social life in starlings. In previous studies, it has been shown that starlings tended to react differently to familiar and less familiar song (Hausberger et al. 1997). Starlings also recognize species-specific songs and react appropriately to them despite local variations in dialect (Adret-Hausberger 1982).

We hypothesized that individuals from more recently established populations in potentially unfamiliar, unpredictable or sub-optimal environments at propagation fronts (Boyd and Richerson 1988) and in habitats disturbed by human activities, such as towns, are more sensitive to social acoustic cues from individuals from other groups than would be the case for individuals from long-established populations. In order to test this hypothesis, we compared the reactions to an acoustic social stimulus between various populations in different colonization phases. Responses of individuals to the playback of a chorus of unfamiliar starling songs were observed in more recently established populations in France and Italy (towns, propagation fronts) and older rural populations in Brittany.

## MATERIAL AND METHODS

### ANIMALS AND SITES

We compared the reactions of starlings from five geographical areas where populations had different colonisation histories (Figure 1):

- Two “old” populations from western France, Brittany, 110 km apart, where starlings became established more than five centuries ago (Richard 1826) and which have abundant food and nesting resources (enough insects and enough available nesting holes) (Clergeau, 1981),

- Rural Rennes: rural population in the country surrounding Rennes city
- Rural St Brieuc: rural population in the country surrounding Saint Brieuc city
- Three “more recent” populations where resources are scarce and less predictable and where starlings have been established for less than 80 years:

- Urban Rennes: 30-80 year old breeding population (Clergeau 1981). This population uses habitats frequently disturbed by anthropogenic activities and with scarce resources particularly during the breeding period (Mennechez and Clergeau 2006).
- Rural Italy: rural populations in the Apuglia region in the recent colonization front established less than 30 years ago (Castiglia and Tabarrini 1982)
- Urban Italy: urban populations of Bari and Mola di Bari cities in Apuglia, also belonging to the colonization front, and established for less than 40 years (Pasquali 1984)

**FIGURE 1:**
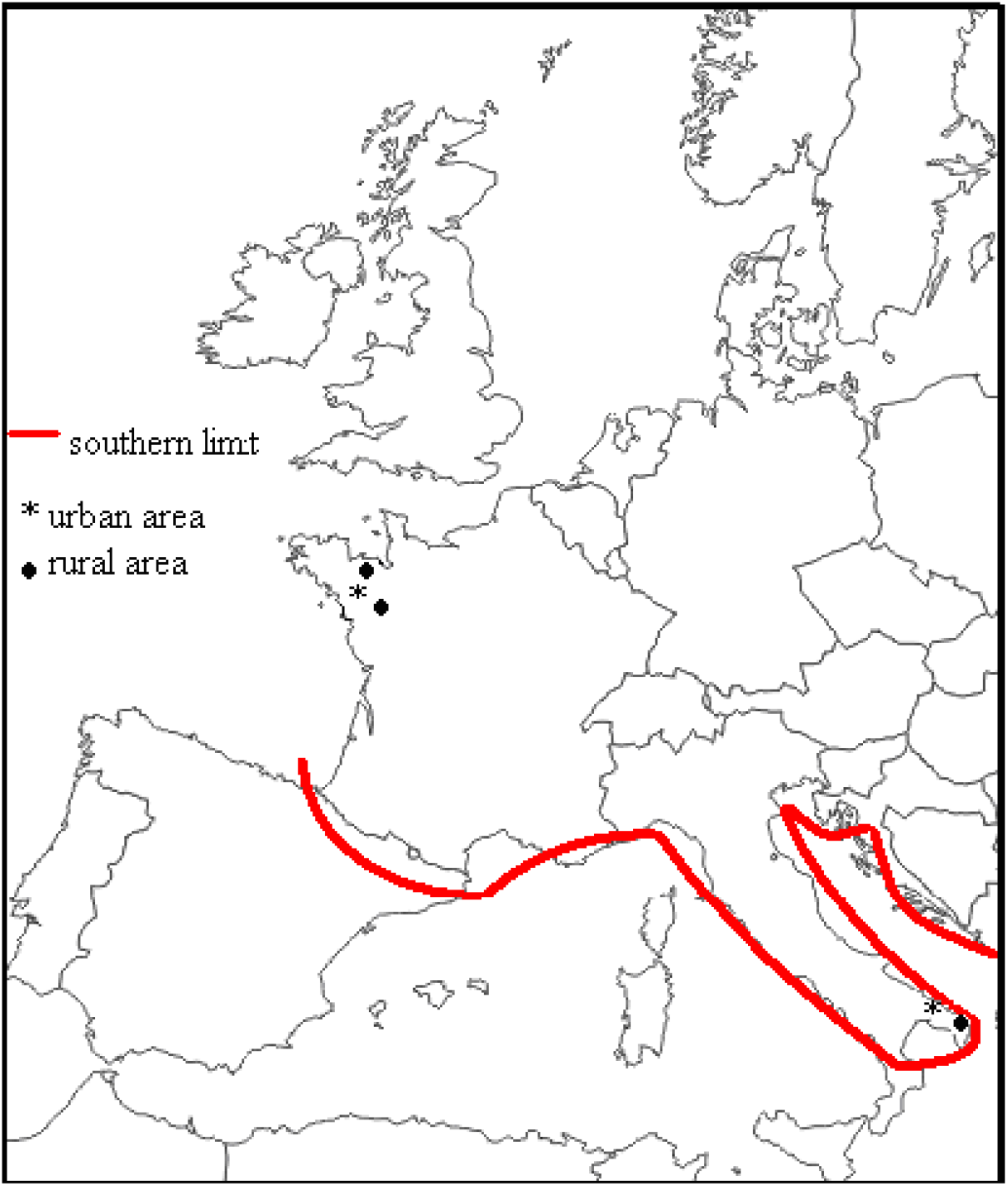
Levels of overall responsiveness to the acoustic stimulus by five populations with different histories

### BEHAVIOURAL TESTS

Selected sites were short grass areas (park lawns, football areas, grazed grass fields) as similar as possible (size between 60m x 45m and 120m x 90m). The study was conducted during the breeding period (April 2008) and during hours of intense foraging and social activity: from 06h30 to12h00 and from 15h00 to 19h00 (Clergeau 1981, Feare 1984). We placed a playback speaker at each site where starlings had been observed and begun playback without preidentified individuals.

In order to make sure that it was the song chorus as such that was attractive rather than local song characteristics, we broadcasted a chorus of distant unfamiliar birds (8 North American starlings: 4 males and 4 females) presenting distant songs. The song structures within the chorus were equally different from both Brittany and Italian local songs (Henry 1994, 1998) and they contained only songs of class II and III (Hausberger 1997) constituting whistles and warbling that do not provide dialectal information. The chorus consisted of continuous singing for 3 min 26 that was repeated for the duration of the test (30 min) between one display and the follower there were only two seconds of pause. This procedure allowed flying birds to randomly hear several different whistles and warbling songs as they could arrive in the observation area at any moment during the chorus. The intensity of playback diffusion was 85 dB when measured at 1m from the loudspeaker. The loudspeaker was placed on the ground one meter from a tree or an electric line in order to allow birds to perch nearby, and it was hidden under a khaki coat. The observer (A.R.) remained 20m away from the loudspeaker and observed and noted the number and behaviour of starlings flying over the observation area (2500m^2^ around the loudspeaker). We chose to test starlings on ten to fifteen sites in each area rather than to focus on only one group per population and we could thus take into account the distribution and density of the colonies. Only one test was performed per site. More than 700 birds were sampled (100 to 180 per population) (Table 1).

**TABLE 1:**
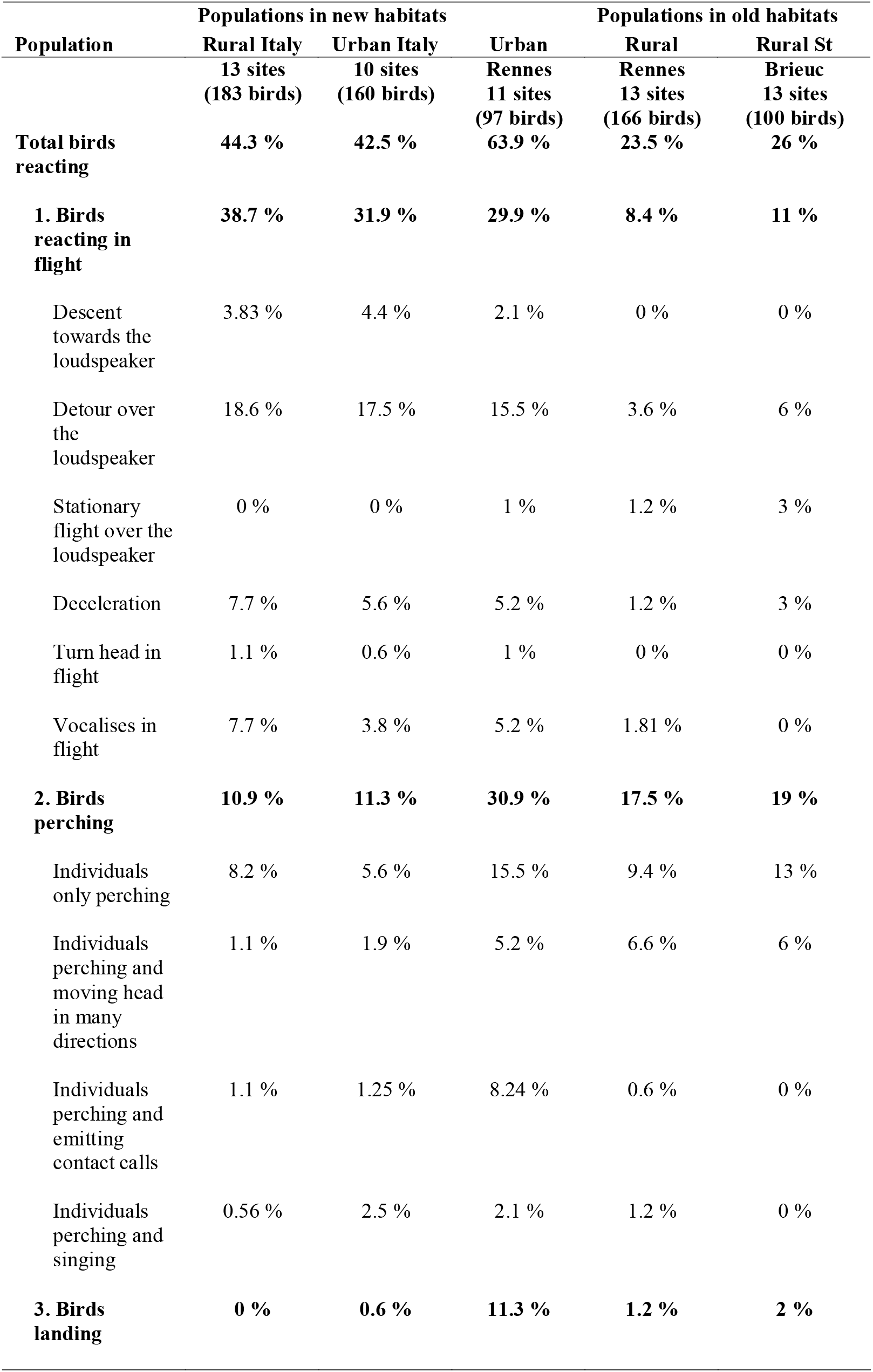
Percentages of individuals that reacted with each kind of behaviour to the playback () = number of birds tested for each population - = not enough individuals to conduct chi^2^ analysis *** p<0.001 ** p<0.01 * p<0.05 df = degrees of freedom

### BEHAVIOURAL RESPONSES

Previous studies on behavioural responses of birds to playback experiments had identified different degrees of reaction. In studies of parent recognition by penguin chicks, the authors distinguished various responses that differed in strength (Aubin and Jouventin 1998, Jouventin et al. 1999). Following this method, we classified the observed behaviours (see Table 1) into 4 categories which could reflect different levels of reactivity:

- no reaction: the bird flew over with no change in direction or speed;
- reactions during flight: the bird could turn its head towards the loudspeaker, or make a detour over the loudspeaker or descend towards it (without landing), or stationary flight over it, or vocalize during flight or even decelerate;
- perching: the bird could simply perch on the tree or electric pole near loudspeaker or perch and turn head searching in many directions, or perch and emit contact calls or sing;
- landing near the loudspeaker.

The experiments were conducted near breeding areas where birds had previously been observed making flights to search food or nest material; they were mainly observed alone or in pairs. In a few cases we observed groups of more than two birds. In any case we considered that a bird was attracted by the sound source only if it was flying and reacting alone or if it was among the first simultaneous birds reacting. We did not consider individuals following their group members as individuals attracted by the chorus in order to avoid an overestimation of the number of reactive birds to the external stimulus. The follower birds were considered as birds who did not react to the chorus. Each observation session lasted 30 minutes. We checked the direction of birds’ flight to avoid recounting the same bird in a same session and only one session was conducted per site in order to avoid recounting the same birds twice. If we were in an area with apparently low bird density and if some birds stayed in the observation area, we stopped the observation earlier to avoid pseudoreplication by this way, we avoid counting the birds more than one time. During the observations, single birds or birds in little groups flew over the experimental area and we counted carefully the number of flying birds and the number of them who reacted to the chorus.

As almost all the birds were flying alone or in groups of 2 or 4 individuals (only four groups of more than 5 individuals) we did not test the effect of group size.

### STATISTICAL ANALYSIS

We have considered the percentage of birds that responded per site. We used pairwise chi square tests (Siegels and Castellon 1988) to compare the differences in acoustic attractiveness between pairs of populations and the differences in response types.

## RESULTS

### OVERALL REACTION TO CHORUS

We observed substantial differences in how birds responded to the acoustic stimulus among our different study populations (χ ^2^ = 27.31, df = 4, p<0.001). In Urban Rennes, about 64% of tested birds reacted to the stimulus whereas in Italian populations we observed between 40 and 45 % of birds reacting. Finally, in Rural Brittany populations less than 26% of the birds expressed a reaction to the playback (Table 1).

In pair-wise comparisons, birds in Rural Italy appeared to be significantly more reactive than those from Rural Rennes and Rural St Brieuc (respectively χ^2^ = 8.24, df = 1, p<0.01 and χ^2^ = 4.32, df = 1, p<0.05). Similarly, the population from Urban Italy showed more reactions than the population from Rural Brittany but this difference was significant only in the pairwise comparison with the Rural Rennes, not with the Rural St Brieuc (respectively χ ^2^ = 6.77, df = 1, p<0.01 and χ ^2^ = 3.51, ns).The Urban Rennes population appeared to be significantly more reactive than the Rural Rennes and the Rural St Brieuc ones (respectively χ ^2^ = 17.81, df = 1, p<0.001 and χ ^2^ = 11.1, df = 1, p<0.001). Finally, the Rural Rennes population and the Rural St Brieuc population appeared to behave similarly (χ ^2^ = 0.13 ns).

In conclusion, more recently established populations appeared to be more reactive towards the playbacks than long-settled ones (Figure 1).

As a corollary of these observations, we observed a gradient of “No reaction” response with three levels:

- the Rural Brittany populations are those that present the highest percentages for this category (around 75% of no reaction).
- the Italian populations with an intermediate level: around 56 % of no reaction
- the Urban Rennes population with a lower absence of reaction: 36 % of no reaction

#### REACTIONS IN FLIGHT

Flight reactions differed across populations (χ ^2^ = 47.25, df = 4, p<0.001) (Table 1): The Rural Italian population was the most reactive in flight (39% of birds); the Urban Italian and the Urban Rennes populations had intermediate levels of response (near 30% of birds); the Rural Rennes and the Rural St Brieuc populations presented the lowest levels of response (10% of birds). We obtained here a gradient of response corresponding to the gradient of age of the populations, the more recent ones being the more reactive.

Making detours and decelerations over the experimental area were the most frequent behaviours observed in all the populations but they were more frequent in recent populations than in older ones (Figures 3A and B, Table 1).

In the Rural Italian population significant more birds made detours than in Rural Rennes and in Rural St Brieuc (χ ^2^ = 15.43 df =1 p<0.001 and χ^2^ = 6.57 df =1 p<0.02 respectively). Urban Italian birds made also significant more detours than Rural Rennes and Rural St Brieuc (χ ^2^ = 13.66, df = 1, p<0.001 and χ^2^ = 5.65, df =1, p<0.02). Urban Rennes population performed significant more detours than Rural Rennes population (χ^2^ = 9.72, df = 1, p<0.01). The other pair-wise comparisons on detour behaviour were not significant (χ^2^ < 3.74, df = 1, p>0.05).

Concerning decelerations, Rural Italy birds decelerated significantly more frequently than Rural Rennes birds (χ^2^ = 7.57, df = 1, p<0.01) and Urban Italy birds decelerated significantly more frequently than Rural Saint Brieuc birds (χ^2^ = 4.56, df = 1, p<0.05). The other pair-wise comparisons were not significantly (χ^2^ < 2.22, df = 1, p>0.05) or the data were inadequate for statistical testing.

Descent flights towards the loudspeaker and turning the head during flight were rare and only expressed in recently colonized habitats. In addition, we observed more vocalisations during flight in recent populations than in older ones. Italian Rural population (with almost 8% of birds vocalizing in flight) vocalised significantly more than Rural Rennes birds (χ^2^ = 5.84, df = 1, p<0.01) and than Rural St Brieuc birds that never vocalised in flight (χ^2^ = 7.46, df = 1, p<0.01) (figure 2C).

**FIGURE 2:**
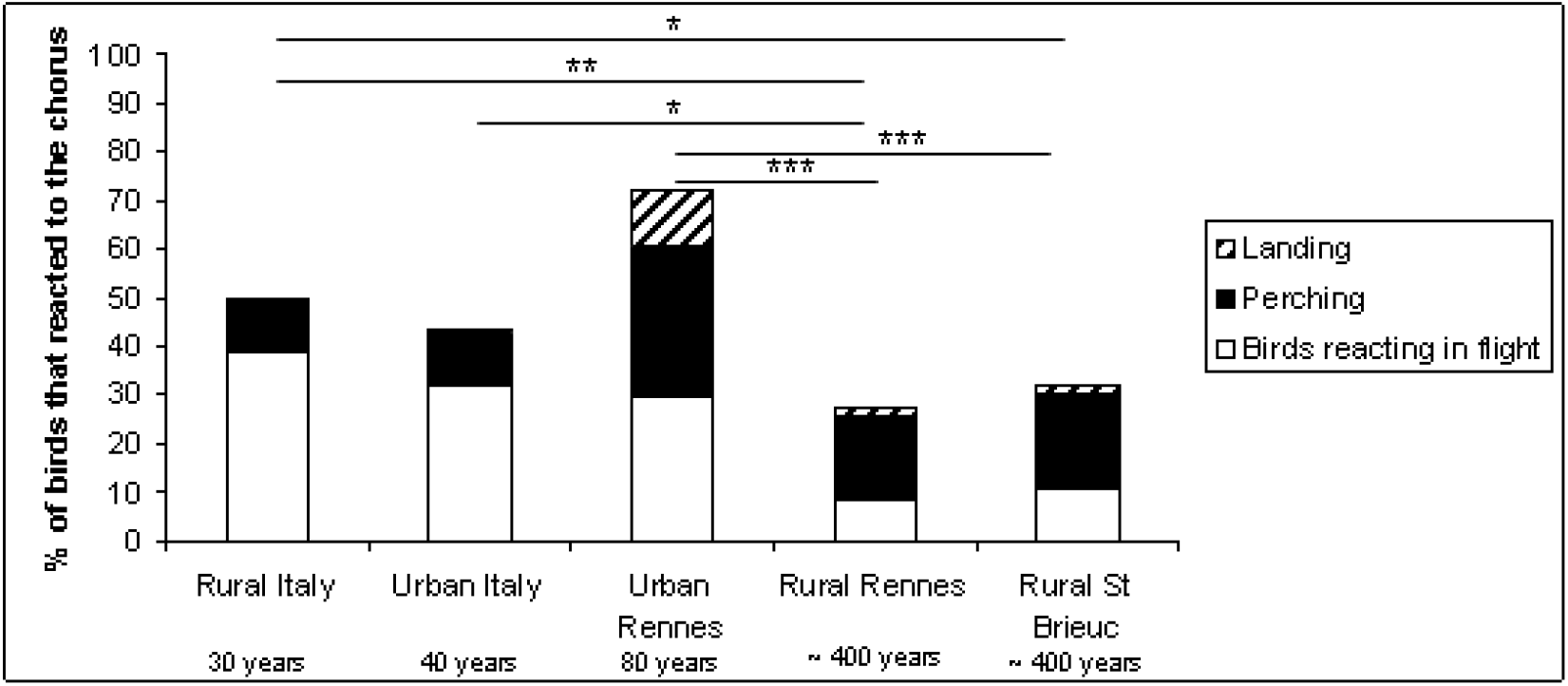
Qualitative and quantitative differences in the expression of responsiveness to the acoustic stimulus. A: Birds making a detour. B: Birds decelerating. C: Birds vocalizing in flight. D: Birds turning the head in many directions when perched. E: Birds singing when perched.

**FIGURE 3.**
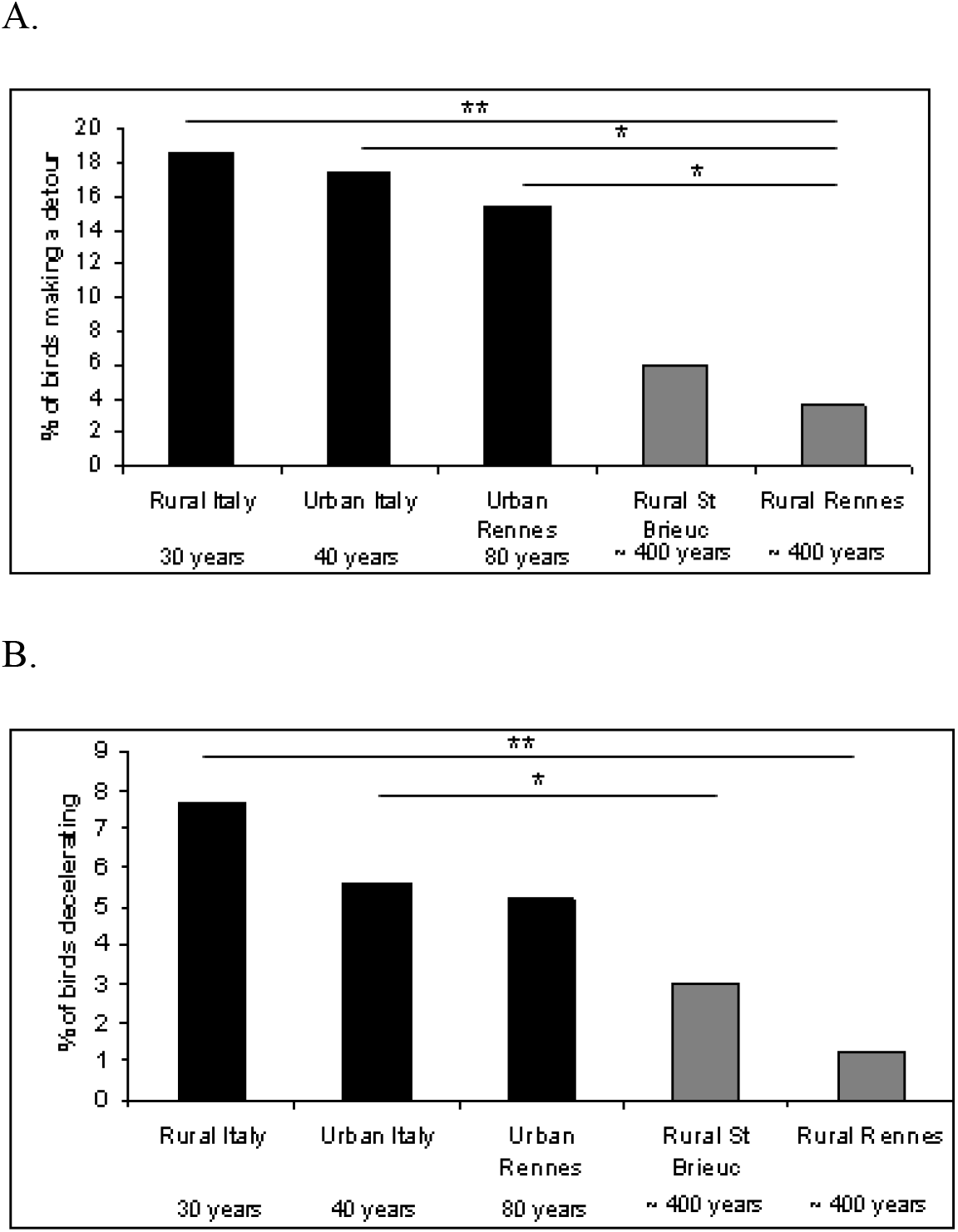
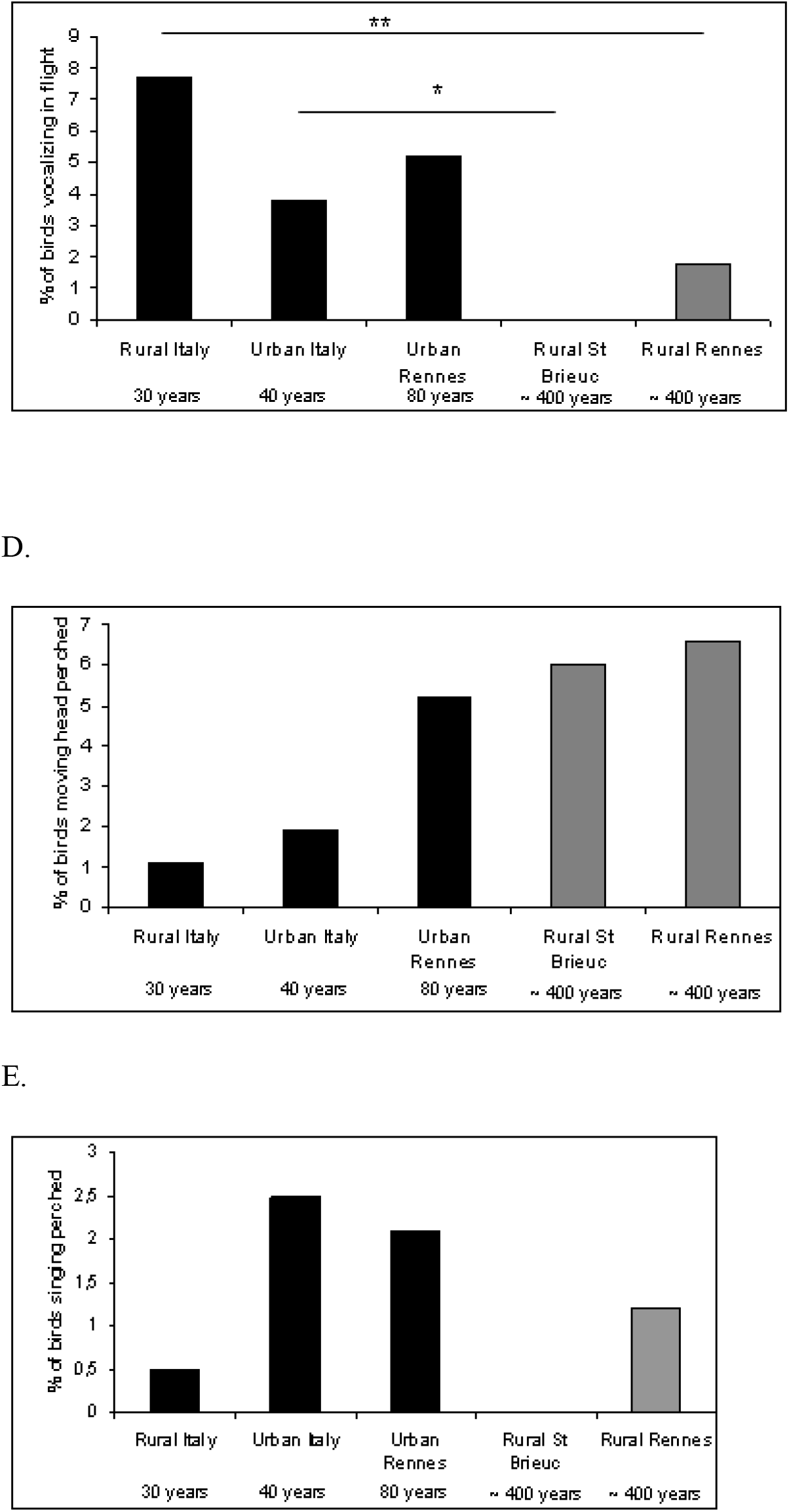

#### PERCHING BEHAVIOURS

Italian populations perched the least (11% of the birds). About 18 % of Rural Brittany birds perched. Urban Rennes population perched the most frequently (31%) (Figure 2).

Some birds, mostly from Brittany populations moved their head in many directions as if they were searching for the singing individuals of the chorus. This behaviour tended to be more frequent in Brittany populations than in Italian ones (Figure 3D) but there were not enough data to conduct pair-wise comparisons. Other individuals vocalized or sang after perching with no differences among populations (Figure 3E).

#### LANDING REACTIONS

Landing on the ground close to the loudspeaker was a rare behaviour and almost only observed in Urban Rennes population (Table 1, Figure 1).

## DISCUSSION

Our experiments show that the studied populations reacted differently to broadcasts of chorus songs. For the first time a higher sensitivity to this kind of social cue was observed in more recent populations than in populations living in long-colonized environments. In a related study, inter-individual differences in sensitivity to social cues were observed by Hahn and Silverman (2006) who observed the reaction of migratory adults of the American redstart (*Setophaga ruticilla*) to song playbacks in order to test the influence of social acoustic cues on habitat selection. They compared experimental plots with playback stimulus and control plots without the acoustic stimulus. They found that experimental plots attracted more adult males that were new immigrants in this geographic area than the control plots, indicating that the settlement in the new habitat by naïve individuals who did not know the area was enhanced by these social cues.

Detecting predators, finding conspecifics, and locating breeding and foraging areas may be crucial when establishing in new environments. Being more reactive to social stimuli could have been an advantage for the ancestors of the more recent populations and could have acted as a mechanism allowing groups’ formation during the settlement process. If this behavioural character was transmitted from the settler generations to the next, the higher sensitivity to social cues observed in recent populations is probably a consequence of this mechanism. It has been shown that invasive ants form large colonies presenting low levels of territoriality that allow proximal gathering between neighbour nests (Holway et al. 1998, Suarez et al. 1999). In our case, gathering with conspecifcs seems to have been a factor allowing colony formation in the past which is still observed today.

Propagation fronts are the expanding edges of a species distribution. The proportion of immigrant individuals can be higher than in long-colonized habitats where on the contrary immigrants are probably less frequent and where the proportion of long time residents can be higher. By consequence, propagation fronts can present higher proportions of naïve individuals that do not know well these areas. Studies on different taxa like bats and birds have shown that naïve, inexperienced or unsuccessful individuals tended to follow successful foraging individuals that had previously located suitable breeding or feeding sites (Wilkinson 1992, Marzluff et al.1996, Kert and Reckardt 2003). In some species as in greater spear-nosed bats (*Phyllostomus hastatus*) vocalizations such as contact calls increase the attractiveness of suitable foraging or roosting sites (Wilkinson and Wenrick Bougman 1998). Sensitivity to social, and in the present case, to acoustic cues is therefore probably an advantage for invasive species.

In their study on responses to shared and non shared song types, Hausberger et al. (1997) found that captive male starlings reacted little to an unfamiliar female song, and that females reacted mostly to song-types they shared with their social partners when these song-types where reproduced by a playback. Here we found that individuals from more recently established populations were very sensitive to the playback even if it was a chorus produced by unfamiliar birds, which means that these starlings react more to the social information even when produced by unknown individuals. Hazlet et al. (2003) had observed this kind of phenomenon at an interspecific scale; they found that two species of introduced invasive crayfish were able to use a broader range of chemical cues than two non invasive species.

Here we interpreted the high levels of response in propagation fronts or in new contexts (like in cities) as the consequence of the recent history of the populations and as a way facilitating colonization. However, the differences of response may be due to differences in food resources or by differences in levels of competition to obtain these resources. Moreover we did not control for ambient noise levels, so this factor could be acting on birds receptivity. Mockford and Marshall (2009) observed that male Great tits from quiet territories reacted significantly stronger when hearing song from another territory holder with low background noise than from those with high background noise. Urban environments are usually noisier than rural ones so birds our birds could have beeen less responsive in urban areas but it was not the case which means that even with noise birds from the recent populations of the urban areas heard well the playback and were more reactive. However more studies controlling both for food resources and background noise should be conducted to confirm if the history of population is really the factor influencing the responses.

Finally, there is an alternative hypothesis to explain the higher sensitivity of birds from recently-established populations. It has been demonstrated that density can influence the use of social acoustic cues (Fletcher 2007). We can thus not exclude the possibility that the higher levels of sensitivity in propagation fronts (southern Italy) are due to the fact that starling density is moderate in this region compared to the higher densities in Rural Brittany. Fletcher (2007) has found that individuals were more sensitive to acoustic cues in habitats with moderate density values. Birds less habituated to hearing unfamiliar songs in areas with low densities during the breeding period may be in consequence more sensitive to social acoustic cues from unfamiliar birds. However this hypothesis can not be applied to the urban Brittany population where bird densities are equivalent to those of the older Rural Brittany population

(Mennechez 1999). In this case the higher levels of reactivity might also be due to the fact that city is a sub-optimal environment where food for nestlings is difficult to find (Mennechez and Clergeau 2006), probably increasing the value of social information.

### BEHAVIOURAL DIFFERENCES IN POPULATION REACTIONS TOWARDS THE CHORUS

Populations differed not only in their overall response but also in their modalities of response. Individuals from more recently established populations reacted mostly by modifying the direction of their flight, perching and vocalising, whereas the major response of older populations was perching with little interaction with the sound source.

In contrast to studies involving visual cues (Clergeau 1981, 1982) where starling decoys were put on the ground, we observed very few landing responses perhaps because the individuals could not visually detect the birds that sang the chorus.

Individuals from more recently established populations tended to vocalise more during flight in response to the acoustic stimulus than individuals from long-established populations. We suggest that individuals may search actively for gatherings of conspecifics and seek out vocal interaction, as they both receive and produce acoustic cues or that calling is contagious as observed in Pampas Meadowlarks (*Sturnella defilippii*) (Gochfeld 1978).

Some individuals moved their head in many directions. Vocal and visual reactions are probably used to obtain additional information and detect and locate the singers. Studies conducted on primates indicated that in the absence of visual obstacles or when inter-individual distances were short, animals used predominantly visual contact to detect and locate conspecifics. On the contrary, when confronted with a habitat containing various visual obstacles (trees or buildings for example) individuals produced contact calls more frequently (Snowdon and Hodun 1981, Koda et al. 2008). In our study flying individuals from Rural Italy tended to vocalize more than individuals from Urban Italy whereas individuals from Urban Rennes vocalized more than those from Rural Brittany. Nevertheless, populations in more recently colonized habitats (either rural or urban) vocalized more than those from older populations, probably looking for more vocal contact. Population history may play a more important role in actively searching for contact than habitat characteristics, but we cannot exclude the possibility that some landscape features could have had an effect on head-turning and vocalising.

These results imply that the use of social cues may act as a major mechanism in social aggregation during colonization or invasion processes and that it is probably a mechanism allowing individuals to join conspecifics in potentially suitable sites (Clergeau 1981, 1982). During establishment in a new area individuals can be confronted with many unknown and novel contexts, new landscapes, new food resources, unpredictable resources, lack of information about predator location, etc. One possible way to compensate for this lack of information is the use of social information gathered from conspecifics, which allows them to settle more easily in the new habitats. Even if other studies are needed to corroborate our results (especially translocation experiments or integration of the levels of inter-individual competition), we can conclude that cues like acoustic signals could play an important role in the colonisation process for gregarious birds such as the European starling.

## ACKNOWLEDGEMENTS

We thank Jean Pierre Richard for his help and explanations of acoustic methods, Patricia Le Quilliec, Giuseppe La Gioia (Museum of Natural History of Salento) and Giovanni Ferrara (University of Bari) for their help in the field. We are grateful to Pr. Adrian Craig for improving the English of the text. Alexandra Rodriguez was supported by a graduate scholarship from the French Ministry of Research and the field experiments were supported by the INRA Scribe Laboratory.

